# Spatial modulation of individual behaviors enables collective decision-making during bacterial group migration

**DOI:** 10.1101/2021.02.17.431709

**Authors:** Yang Bai, Caiyun He, Junjiajia Long, Xuefei Li, Xiongfei Fu

**Affiliations:** Guangdong Provincial Key Laboratory of Synthetic Genomics, CAS Key Laboratory for Quantitative Engineering Biology, Shenzhen Institute of Synthetic Biology, Shenzhen Institutes of Advanced Technology, Chinese Academy of Sciences, Shenzhen, 518055, China; University of Chinese Academy of Sciences, Beijing, 100049, China; Yale University, Department of Physics, New Haven, CT 06520-8120, USA

**Author notes:** Yang BAI & Caiyun HE contributed equally to this work.

## Abstract

Coordination of individuals with diversity often requires sophisticated communications and high-order computational abilities. Microbial populations can exhibit diverse individualistic behaviors and yet can engage in collective migratory patterns with a spatially sorted arrangement of phenotypes following a self-generated attractant gradient. However, it’s unclear how individual bacteria without complex computational abilities can achieve the consistent group performance and determine their positions in the group while facing spatiotemporally dynamic stimuli. Here, we investigate the statistics of bacterial run-and-tumble trajectories during group migration. We discover that, despite of the constant migrating speed as a group, the individual drift velocity exhibits a spatially dependent structure that decreases from the back to the front of the group. The spatial modulation of individual stochastic behaviors constrains cells in the group, ensuring the coherent population movement with ordered patterns of phenotypes. These results reveal a simple computational principle for emergent collective behaviors from heterogeneous individuals.

## Introduction

Collective group migration as an important class of coordinated behaviors is ubiquitous in living systems, such as navigation, foraging, and range expansion (Krause, Ruxton et al., 2002, Partridge, 1982, Sumpter, 2010). In the presence of individual heterogeneity, the migrating group often exhibit spatially ordered arrangements of phenotypes (Krause et al., 2002, Parrish & Edelstein-Keshet, 1999, Partridge, 1982, Sumpter, 2010). In animal group migration, individual behavioral abilities (e.g. directional-sensitive) would result social hierarchy, which further drives the spatial arrangement in coordinated group (Couzin, Krause et al., 2005). At the same time, the spatial arrangements can lead to different costs and benefits for the individuals participating in the group migration (Krause, 1994, Parrish & Edelstein-Keshet, 1999, Partridge, 1982). Participating individuals must follow disciplinary rules to organize themselves into coordinated patterns while on the move, which requires complex computational abilities to interact with the group and the environment (Couzin & Krause, 2003, Couzin, Krausew et al., 2002, Vicsek & Zafeiris, 2012). Therefore, understanding how individuals of different phenotypes determine their group positions is an essential prerequisite to uncover the organization principles of collective populations.

The chemotactic microbe, *E. coli*, provides a simple model to address the emergence of collective decision-making, as it can both exhibit individualistic behaviors (Dufour, Gillet et al., 2016, Frankel, Pontius et al., 2014, Kussell & Leibler, 2005, Waite, Frankel et al., 2016, Waite, Frankel et al., 2018) and collective migratory patterns (Adler, 1966a, Fu, Kato et al., 2018, Keller & Segel, 1971, Wolfe & Berg, 1989). Individual cells perform run-and-tumble random motions by spontaneous switching the rotating direction of flagella (Berg, 2004, Berg & Brown, 1972). The cell can facilitate the chemotaxis pathway to control the switching frequency to bias in its favorable directions towards the chemoattractant gradient, where the efficiency to climb the gradient is defined as chemotactic ability (*χ*) (Celani & Vergassola, 2010, de Gennes, 2004, Dufour, Fu et al., 2014, Si, Wu et al., 2012). In addition, substantial phenotypic heterogeneity in chemotactic ability has been observed even for clonal bacterial population, which diversifies the chemotactic response of cells to identical signals (Spudich & Koshland, 1976, Waite et al., 2016, Waite et al., 2018). At the same time, despite of the stochastic solitary behavior and variations in phenotypic ability, *E. coli* population can migrate as a coherent group by following a self-generated attractant gradient (via consumption) (Adler, 1966a, Saragosti, Calvez et al., 2011, Wolfe & Berg, 1989). The group moving at a constant speed form a stable pattern of phenotypes sorted by their chemotactic abilities (Fig 1A), so as to maintain phenotypic diversity in the coherent migratory group (Fu et al., 2018, Waite et al., 2018). Intriguingly, it’s believed that there are no direct communications among cells within such coordinated migration group (Cremer, Honda et al., 2019, Fu et al., 2018), and cells encounter highly dynamic external stimulus. How individuals with phenotypic and behavioral variations manage to maintain the consistent group performance and determine their relative positions in the group is still a mystery.

**Figure 1.**
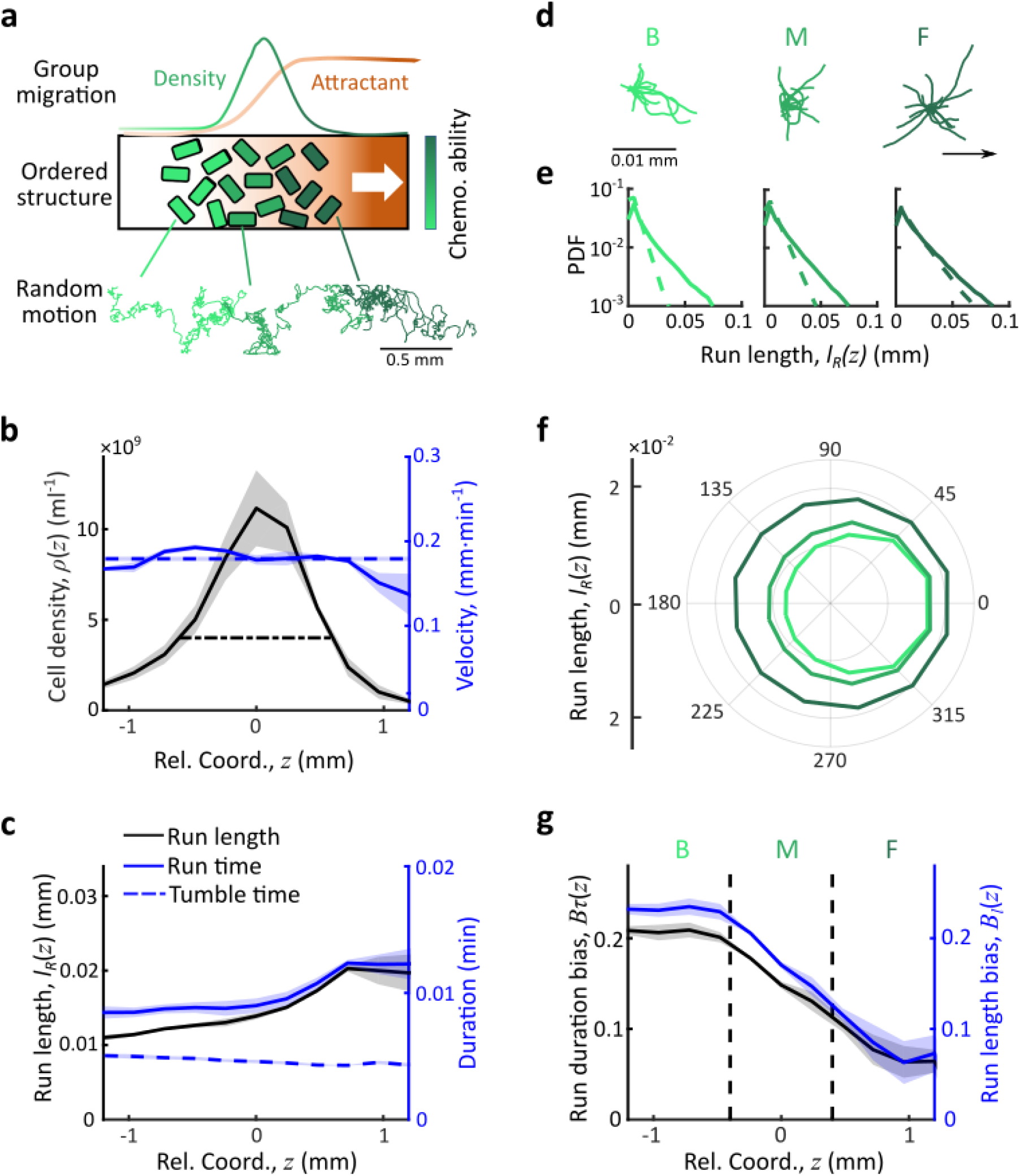
Behavioral structure of individual bacteria in collective group migration. (**A**) Illustration of bacterial chemotactic group migration. Bacteria may form collective migrating group (green line) while consuming chemo-attractant collectively (brown line). Bacterial population of diverse phenotypes are sorted by their chemotactic abilities (increasing from light green to dark green) during collective migration following the self-generated attractant gradient (brown color). Meanwhile, as shown in the sample trajectories, individual cells perform run-and-tumble random motions biased towards the group migration. (**B**) In the moving coordinate *z* = *x* − *V*_*G*_ *t*, the bacteria density profile *ρ*(*z*) is stable (black solid line). The width of the density profile is defined as 2 times the standard deviation of bacterial relative position (2σ, black dash line), represented by the black dash-dotted line. The instantaneous velocity (*V*_*I*_ (*z*)) (blue solid line) is uniform and equals to the average group velocity *V*_*G*_ (blue dash line). (**C**) The mean run length ⟨*l*_*R*_(*z*)⟩ (black solid line) and run time ⟨*τ*_*R*_(*z*)⟩ (blue solid line) increase from the back (left) to front (right) of the migration group, while the mean tumble time ⟨*τ*_*T*_(*z*)⟩ slightly decreases (blue dash line). (**D**) Sample runs of bacteria from the 3 regions. B, M, and F stands for the back, middle and front of the migration group, respectively. Regions were defined by black dashed lines in *G*. Trajectories of runs show that cells in the back of group tend to run forward, compared to cells in other regions. (**E**) The exponential distribution of forward run length (solid lines) and backward run length (dash lines) in 3 regions show that the difference of run length between forward and backward for cells in the back is larger than cells in other regions. (**F**) The mean run length in different directions, with the angular bin size of 15°, also show that cells in the back were better skewed to run forward. (**G**) The run length bias 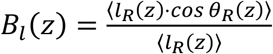 (black solid line) and the run time bias 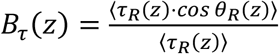 (blue solid line) both decreased from the back to the front of the migration group, which is also consistent with results shown in D-F. In panel B, C, G, shaded area represents s.e.m. of 3 biological replicates. The spatial bin size is 240*μm*.

Here we analyzed bacterial trajectories in the chemotactic group migration using a microfluidic system that enables us to simultaneously characterize the quantitative properties of individual motions (see Materials and Methods). We discovered that in the collectively migratory group, the run-and-tumble motions of individual cells were spatially modulated to behave as mean-reverting processes relative to the group, i.e. cells effectively tend to revert its direction of runs towards the mean position of the group. The same rule of behavioral modulation applies to cells of different phenotypes to allow them migrate at a consistent average speed with an ordered spatial arrangement of phenotypes. By titrating the phenotypes with different chemo-receptor abundance, we further demonstrated that the mean-reversion rate of the behavioral modulation depends on the sensitivity response to the chemoattractant gradient. Therefore, although the high-order computational abilities are not available to the simple organisms, the spatial modulation of stochastic behaviors at the individual level enables novel decision-making capabilities at the population level.

## Results

### Spatially ordered bacteria behavior

To directly investigate how bacteria with different chemotactic abilities determine their relative positions within the collective migration group via run-and-tumble random motions, we employed a Y-shape microfluidic device with a long channel of 20mm which allows us to generate a stable propagating band of bacteria as previously reported (Fu et al., 2018, Saragosti et al., 2011). Specifically, about 1.5 × 10^4^ *E. coli* wild type cells (strain RP437) were loaded into the device, and the medium used is M9 motility buffer supplemented with 200μM aspartate (Asp) as the only chemo-attractant in the system (Adler, 1966b, Fu et al., 2018). Under this condition, only one dense band of migrating bacteria can be spontaneously formed (Fu et al., 2018), after the cells were centrifuged to the tip of the long channel. To quantify the statistics of the single-cell motions within the dense traveling band, we premixed a small fraction of bacteria (strain JCY1) which constitutively expresses yellow fluorescent protein with the non-fluorescent wild type population (strain RP437) by 1:400. The trajectories of fluorescent cells were then recorded under 4X objective with a frame rate of 9 fps for 10 mins (see Materials and Methods and Fig S1). As the fluorescent labeled cells show the same behavior as the wild type ones (Fig S1D), we can consider the behavior of fluorescent cells as the representatives in the migrating group (Fu et al., 2018, Saragosti et al., 2011).

As an important advantage of the experimental setup, we can trace the single cell motions within the dense band of group migration for long time (e.g. typical tracks are larger than 300 seconds) (Saragosti et al., 2011). Given the long trajectories of fluorescent bacteria, we first observed an overall trend of biased random motions of individual cells towards the group migration (Movie. S1). By analyzing the instantaneous velocity based on the trajectories of fluorescent bacteria projected to the group migration direction *x*_*i*_(*t*), we found that the average instantaneous velocity of the entire group, *V*_*G*_(*t*) = ⟨*Δx*_*i*_(*t*)/*Δt*⟩, kept constant over time, *V*_*G*_ ~3.0 *μm*/*s*, suggesting that the band of cells as a whole propagates at a constant speed (Fig S2). Consequently, in the moving coordinate (*z* = *x* − *V*_*G*_ *t*), the time-shifted cell density profiles *ρ*(*z*) can be superimposed as an approximately invariant profile (Fig 1B). Furthermore, the average instantaneous velocity over the band profile, *V*_*I*_ (*z*) = ⟨*Δx*_*i*_(*z*)/*Δt*⟩, remained the same as the group velocity (*V*_*G*_ = ⟨*V*_*G*_ (*t*)⟩_*t*_) along the density profile (Fig 1B). We also verified that these observations did not depend on the sampling time interval (Fig S2). Therefore, despite of stochastic motions on single cell level, bacterial population are able to migrate as a stable group (Adler, 1966a, Fu et al., 2018, Saragosti et al., 2011).

Next, to address how the collective group movement emerge from the stochastic solitary behavior, we analyzed the statistics of run-and-tumble events for individual bacteria. Specifically, after identifying all the run states of individual trajectories by a previously described computer assistant program (see Materials and Methods) (Dufour et al., 2016, Waite et al., 2016), we aligned all the time-shifted runs by their starting positions in the moving coordinate with respect to the center of the group, and quantified the spatial distributions of the mean run duration ⟨*τ*_*R*_(*z*)⟩, as well as the mean run length ⟨*l*_*R*_(*z*)⟩. We find that both ⟨*τ*_*R*_(*z*)⟩ and ⟨*l*_*R*_(*z*)⟩ increase from the back to the front of the moving group, while the mean tumble time ⟨*τ*_*T*_(*z*)⟩ is almost invariant (Fig 1C). Note that previous observations on the steady state profiles of population distribution exhibited that phenotypes with low tumble bias would spontaneously position themselves in the front of the migratory group (Fu et al., 2018). Besides the phenotypic distribution along the wave profile, the spatial structure of run length/duration can also be contributed by the modulation of bacterial behaviors in response to the chemoattractant gradient (Dufour et al., 2014, Long, Zucker et al., 2017, Shimizu, Tu et al., 2010). It’s unclear how the individual behaviors are dynamically modulated during the group migration.

To answer this question, we first investigated sample runs initiated from the back (B), middle (M) and front (F) of the migration group. Qualitatively, the lengths of representative runs in the front are longer but distribute more uniformly in terms of the directionality, whereas the lengths of runs in the back are shorter but the directions of runs are more likely pointing towards the direction of group migration (Fig 1D). Quantitatively, the statistics of run lengths, as well as the run durations, display exponential distributions, suggesting that the switch between runs and tumbles follow Poisson process (Berg & Brown, 1972, Wang, Shi et al., 2017). Of those distributions, the means in the direction of group migration are longer than that of the opposite direction (Fig 1E and Fig S3A). Furthermore, by analyzing the angular distribution of the run length, we found that in the front of the group, the difference between the forward runs and backward runs became smaller despite increased mean values (Fig 1F). Moreover, we also observed that the reorientation angles after tumble events exhibited a decreasing trend along the wave profile (Fig S3E), suggesting a directional persistence towards the group migration as previously reported (Saragosti et al., 2011). All these results suggest that the bacteria in the back run more effectively towards the group migration than those in the front.

To further quantify the efficiency of runs, we calculated the directional bias of run length and run duration, which are defined as the ratio of the net run length/duration in the direction of the group migration and run lengths/durations in all directions, 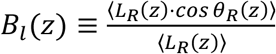, and 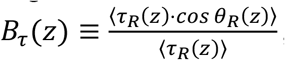, respectively, where *θ*_*R*_ is the angle between single runs and the migration direction. Both quantities are spatially modulated as they decreased by 3∼4 folds from the back to the front of the migration group (Fig 1G), quantitatively indicating how much more effectively the cells in the back behave than that in the front. As the tumble duration is almost constant along the band profile (Fig 1C), we hypothesized that the efficiency of runs would represent how fast that the cells climb the chemoattractant gradient, suggesting that the spatial modulation in the directional bias of runs enables cells in the back of the migration group to exhibit higher drift velocity though the mean run length of them is shorter.

### Bacteria perform mean-reversion behavior as active particles in moving gradient

To verify our hypothesis, we examined the expected drift velocity along the band profile, which is defined as the average projection of run length on the migration direction over the average duration of runs and tumbles, 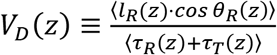 (Dufour et al., 2014). Unlike the spatially-uniform instantaneous velocity *V*_*I*_(*z*), the expected drift velocity *V*_*D*_(*z*) decreases linearly from the back to the front of the migration group with a fitted linear slope of −*r* = −0.05 min^−1^ (Fig 2A). The negative slope of *V*_*D*_(*z*) suggests a mean-reversion behavior of bacteria: the bacteria in the back of the migration group are expected to drift faster than the group (*V*_*D*_ > *V*_*G*_), enabling the cells to catch up within the group, while the cells in the front are expected drift slower than the group (*V*_*D*_ < *V*_*G*_), making the cells to slow down and fall back (Fig 2B). As another piece of evidence supporting the mean-reversion behavior, the time-shifted trajectories of cells relative to the group indicate that the cell motions perform sub-diffusive (Fig 2C), of which the mean square displacement (MSD) are constrained over time (Fig S3G). Thus, our observations indicate that the modulation of individual runs along the wave profile leads to the spatially-structured expected drift velocity, resulting an effective mean reversion process of cell motions.

**Figure 2.**
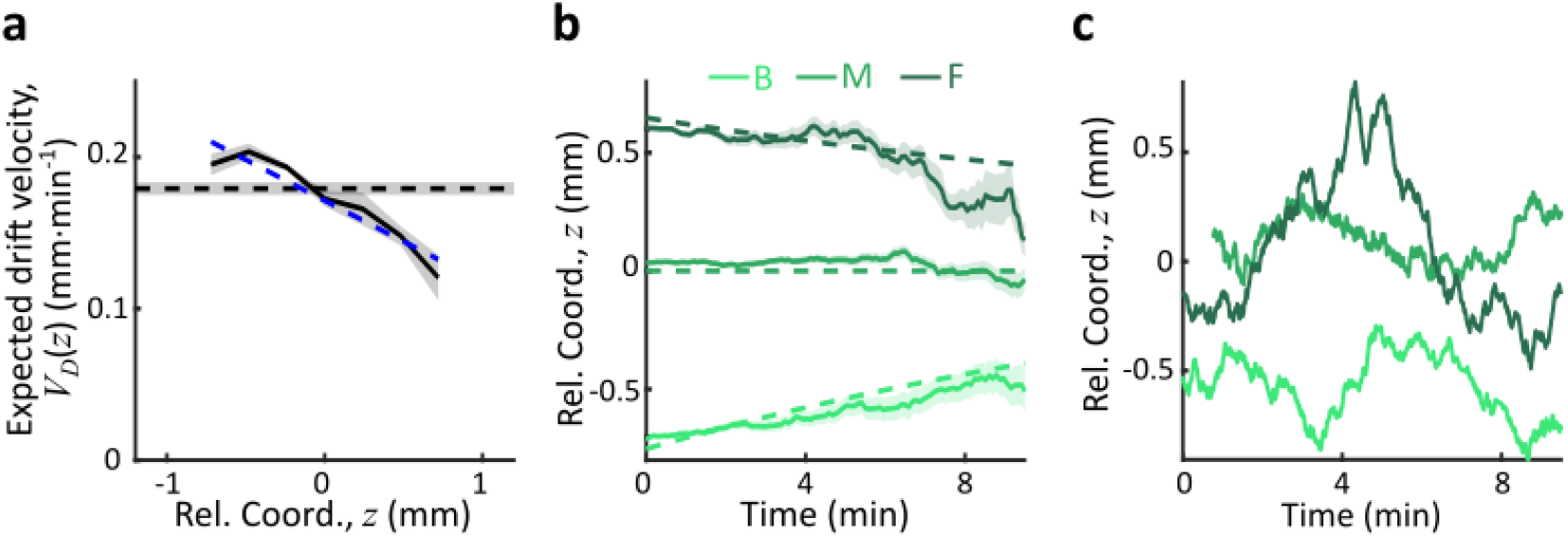
Mean-reversion behavior of individual bacteria relative to the group. (**A**) The expected drift velocity 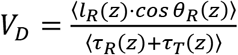 (black solid line) decreases from the back to the front of the migration group. The blue dash line is the linear fit of the quantified experimental data, i.e., *V*_*D*_ ≈ −*rz* + *V*_*D*0_, with *r* = 0.05*min*^−1^ and *V*_*D*0_ = 0.17*mm* · *min*^−1^. *V*_*D*_ crosses with the average group velocity *V*_*G*_ (black dash line), which implies that bacteria perform mean-reversion motions, i.e. cells in the back catch up the group while cells in the front lay back. The *V*_*D*_ curve and its linear fit was cut to present ∼90% majority of cells (±1.65*σ*). (**B**) The time evolution of the average expected position (*z*, solid lines) of cells starting from the back (light green), middle (green), and front (dark green) of the migration group (defined in Fig1. G). Shaded area represents s.e.m. of more than 450 cells (see Methods). Analytically, the O-U type model predicts that *z* = *C*_0_*e*^−*rt*^ − (*V*_*D*0_ − *V*_*G*_)/*r* (dash lines), where *C*_0_ can be fitted by the starting position (see Supplementary text). (**C**) Representative examples of single-cell trajectories (3 colors represent 3 different tracks) showed the reversion behavior of bacteria around their mean positions.

To better understand how the spatial modulation of expected drift velocity emerges, we adopted a one-dimensional minimal model of bacterial behavior. The biased random motions of individual cells are described as an active Brownian particle in the low Reynolds number regime in the medium, following a Langevin type equation *dx*_*i*_ = *V*_*D,i*_*dt* + *𝜖dW* (Berg, 2004). In the stochastic velocity is modelled by a Gaussian random force ϵdW of which the variance ϵ depends on the effective diffusion coefficient of the cells (Rosen, 1973, Rosen, 1974), while the deterministic velocity is the expected drifted velocity *V*_*D*_ = *χg*(*x, t*) that depends on two key parameters (Celani & Vergassola, 2010, de Gennes, 2004, Dufour et al., 2014, Si et al., 2012): the cell chemotactic ability, *χ*, and the perceived attractant gradient *g*(*x, t*) (Eq. S3). To calculate the perceived attractant gradient, we considered the dynamics of attractant concentration *S*(*x, t*) as a diffusible small molecule that can be consumed by the cells (Eq. S2). Such stochastic description is equivalent to the classic Keller-Segel model (Keller & Segel, 1971, Rosen, 1973). For simplicity, we considered each cell has the same attractant consumption rate independent to the local cell density, and omitted the hydrodynamic forces and physical interactions among cells (Drescher, Dunkel et al., 2011, Fu et al., 2018, Saragosti et al., 2011). Using this particle-based model, 100,000 cells were simulated in one dimension. Starting with all cells in one end (*x*_*i*_ = 0) and homogenously distributed attractant field (*S*(*x*) = *S*_0_), a stable band of cells would spontaneously emerge by following a moving gradient of attractant that is generated by cell consumption for both single phenotype and multi-phenotypes (Fig S4). In the presence of diversity in cell chemotactic ability, the traveling band exhibits a sorted structure of phenotypes as previously observed (Fu et al., 2018). The self-generated perceived attractant gradient in the moving coordinate, *g*(*z*), exhibits a stable profile that decreases from back to front.

In the moving coordinate, the Langevin type equation writes: *dz*_*i*_ = *V*_*D,i*_*dt* − *V*_*G*_ *dt* + *𝜖dW*. It tells us that the motion of each individual cell relative to the group migration can be considered as an active particle regulated by two ‘effective forces’: one generated by the decreasing trend of *V*_*D*_ which pushes the cell to catch up the wave; and another generated by the moving gradient of *V*_*G*_ which leaves the cell fall behind the wave. This mechanism constrains the random motions of cells, and enable cells with different phenotypes to form the spatially ordered structure spontaneously. Specifically, for cells with chemotactic ability *χ*_i_, the balance between the two ‘effective forces’ produces an effective potential well 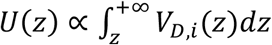 (Eq. S17).

The simulation results (Fig S4F) show that the perceived gradient *g*(*z*) is almost linear along the band profile. Thus, we further approximated *g*(*z*) as a linear function of z around the peak position: *g*(*z*) ≈ *g*_0_ + *g*_1_*z* with *g*_1_ < 0, which gives us an analogy that cells follow an Ornstein-Uhlenbeck (OU) type process in the moving coordinate *dz*_*i*_ = *χ*_*i*_*g*_1_*zdt* + (*χ*_*i*_*g*_0_ − *V*_*G*_)*dt* + *𝜖dW*. Solving this equation, we obtained that cells perform mean reversion motions around the mean positions 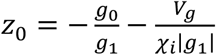 with the reversion rate 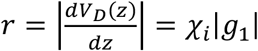 (Supplementary text). As a result, the run-and-tumble random motions of cells are constrained in the potential well, of which the minimum (the same as the mean position of cells, *z*_0_) increases with the chemotactic ability of the cells *χ*_*i*_. In addition, the standard deviation (*σ*) of spatial distributions of cells, given by 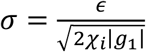, decreases with *χ*_i_.

This analysis suggests that the spatial ordering of cells does not care how is the perceived moving gradient *g*(*z*) is generated, as long as the slope of it *g*_1_ < 0 is negative. Thus, we deduced a non-consumable moving attractant field (*S*(*z*)) from the measured density profile *ρ*(*z*) (Eq. S12), and simulated the behavior of cells following the Langevin type equation under this moving attractant field. As shown in Fig S5, cells with large enough *χ* follows the moving attractant field and are spatially sorted as predicted by the OU type model.

### Ordered effective potential wells for bacteria of different phenotypes

To consolidate the proposed mechanism underlying the emergence of spatial orders from the individual random motions, we further performed simulations for cells of various chemotactic abilities integrated with the chemotactic pathway and multi-flagella competition. Together with the attractant dynamics *S*(*x, t*) described in Eq. S2, we performed stochastic simulations in three dimensions of a population with different chemotactic abilities *χ*_*i*_, where *χ*_*i*_ was varied by tuning the receptor gain *N* (For details in Supplementary text)(Dufour et al., 2014, Jiang, Qi et al., 2010, Sneddon, Pontius et al., 2012). As the receptor gain only affects the amplification factor that a cell responds to the gradient, the variation of bacterial motility *𝜖* is unchanged. As a result, a dense band of migrating cells that follow a self-generated moving attractant chemoattractant gradient via consumption were recaptured as experiments (Fig S6). To better analysis the simulations, we then simplified the model by the assumption of a non-consumable attractant profile *S*(*z*) moving along the group migration direction (Fig S5A). Using this simplified model, we first checked that mean positions of density profiles of cells with different receptor gain *N*, as well as their peaks, were orderly aligned in respect to chemotactic ability *χ*_*i*_.

As an important advantage of the agent-based simulations, the model allows us to analyze the single cell behavior during the ordered group migrations. For each phenotype *i*, the expected drift velocity *V*_*D,i*_(*z*) decreases along the density profile (Fig 3A). Consistent with the ordered structure of density profiles, the intersection between *V*_*D,i*_(*z*) and *V*_*G*_ exhibits the same sorted order of chemotactic ability *χ*_*i*_ (Fig S7). As the reversion rate 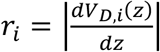 shows a positive correlation to *χ* _*i*_, cells with lower receptor gain *N* (resulting smaller *χ*) experience a weaker reverting force towards centers (Fig 3B). Thus, the effective moving potential, *Ui*(*z*), which constrains the cells round mean positions sorted by their chemotactic abilities, becomes flat for cells with lower chemotaxis ability *χ* (Fig 3C) (Long, 2019). As a result, cells of each phenotype perform as sub-diffusion, of which the MSD along the migration coordinate relative to the group are bounded at the level negatively correlated to *χ* (Fig 3D). We further obtain similar results for populations of different *χ*_*i*_ through adaptation time *τ*, or basal CheY protein level *Y*_*p*0_ which determines the basic tumble bias *TB*_0_ (Dufour et al., 2014, Jiang et al., 2010, Sneddon et al., 2012) (Fig S8).

**Figure 3.**
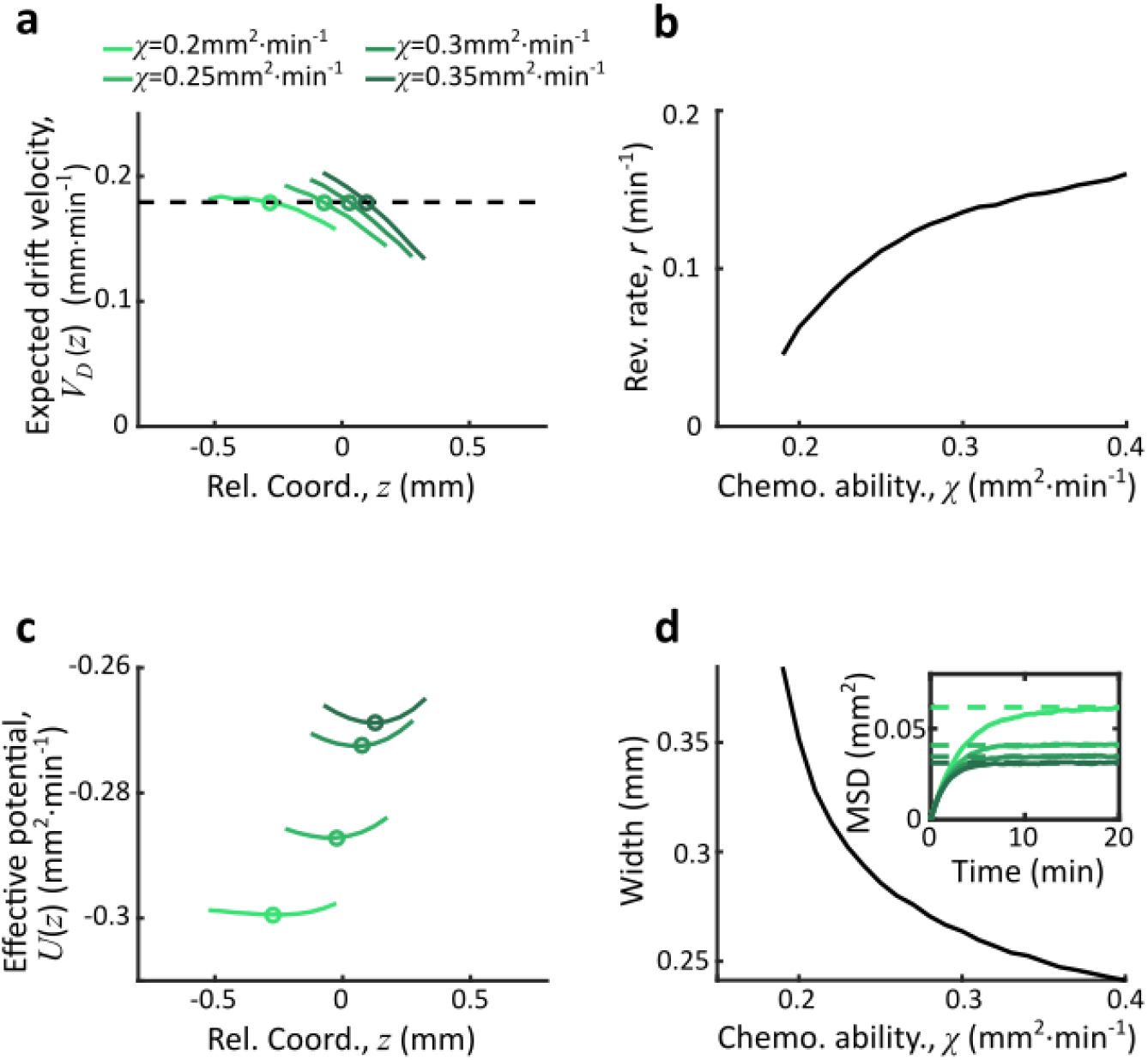
Agent-based simulations recapture the ordered structure of bacterial motions. (**A**) The expected drift velocity *V*_*D*_(*z*) of simulated bacteria decreases from the back to the front of the migration group, where the chemotactic ability *χ* ranges from 0.2 to 0.35 *mm*^2^ · *min*^−1^, consistent with the experimental results shown in Fig. 2a. The intersections between *V*_*D*_ curves with the preset group velocity *V*_*G*_ (black dashed line) shifts towards the back of the migration group as *χ* decreases (circles). Different colors of the lines and circles correspond to different chemotactic abilities *χ* as shown in the legend. The same color-coding also applies to (**B-D**). (**B**) The reversion rate *r*_*i*_= |*dV*_*D,i*_(*z*)/*dz*| increases with the chemotactic ability. (**C**) The effective potential well calculated by 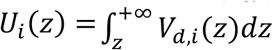. Positions of the potential minimum *z _min_* are marked as circles. As illustrated, for a lower chemotaxis ability *χ*, the potential well is shallower and z_min_ shifts towards the back part of the migration group. (**D**) The width of the density profile (measured by 2*σ*, see Fig. 1B) decreases with the reversion rate r_i_ as well as the chemotaxis ability *χ*_i_. The mean square displacement (MSD) of bacteria (insert, solid lines) is bounded to 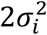 (insert, dash lines) (see Supplementary text). In panel (**A, C**), curves were cut to present 90% majority of cells (*z*_*min*_ ± 1.65*σ*_*i*_). More details of this simulation results were presented in Fig. S7.

To verify the model predictions on the individual behavior of different phenotypes, we experimentally measured the trajectories of cells with different chemotactic abilities during the group migration. Specifically, we altered the chemotaxis abilities of cells by titrating the expression level of Tar, which is under the control of a small molecule inducer aTc (Sourjik & Berg, 2004, Zheng, Ho et al., 2016) (see Materials and Methods and Fig 4A). The variations on the expression of Tar would lead different receptor gains in response to the Asp gradient (Adler, 1966b, Adler, 1969, Tu, 2013), but the tumble bias and growth rate will not change (Fig S9). The tar-titrated cells labeled with yellow fluorescent protein (strain JCY20), were added into wild type population by the ratio of 1 in 400. Within the wild type population, 1 in 50 cells were labeled with red fluorescent protein (strain JCY2). As the tar-titrated strain is a small portion of the pre-mixed population, we can consider the density profile of the population is invariant to different inductions of tar. The premixed population can generate a collective group migration as the wild type population does (Fig S9). The trajectories of YFP labeled cells were tracked to represent the behavior of cells with different chemotactic abilities, while the profile of wild type cells with RFP was also measured to characterize the density distribution of the entire migratory population.

**Figure 4.**
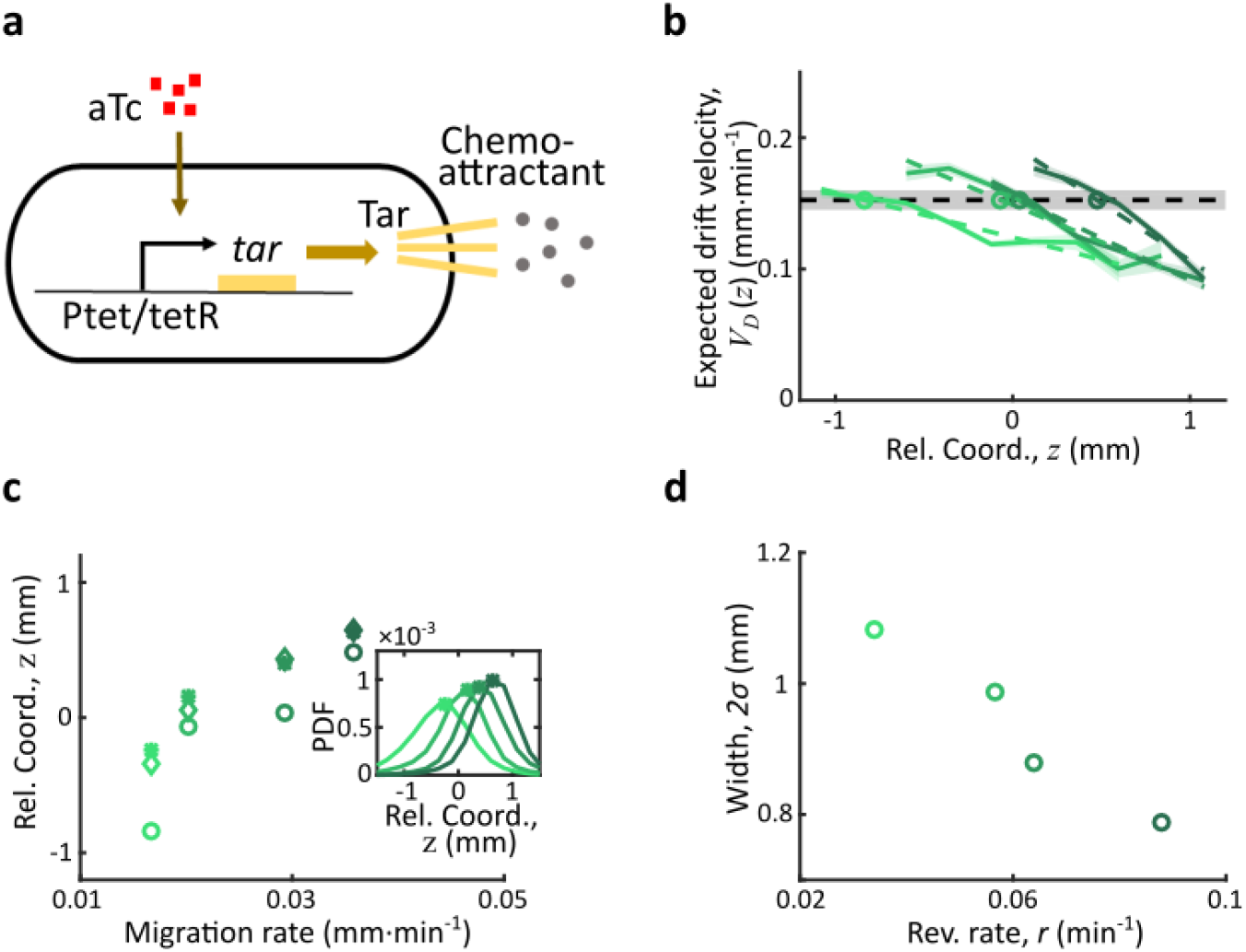
Spatial ordered structure emerged from behavioral modulation of cells with different chemoreceptors. (**A**) Genetic circuit of the Tar titratable strain. In the experiments, the expression level of Tar (a chemo-attractant receptor protein) was titrated by the concentration of external inducer (aTc). The chemotactic ability *χ* of bacteria is then determined by the expression level of Tar (*41*). (**B**) The expected drift velocity *V*_*D,i*_(*Z*) of Tar titratable strain JCY20 (colored solid line) were spatially modulated. All of them decrease from the back to the front of the migration group and intersect with the group migration velocity *V*_*G*_ ≈ 0.15*mm*/*min* (black dash line). The linear fits of *V*_*D,i*_(*z*) (colored dash lines) intersect with V_G_ at positions (circles) determined by the corresponding Tar expression level. Colors from dark to light green corresponds to inducer (aTc) concentration to be [1, 3, 6, 20] *ng*/ *mL*. The black shaded area of *V*_*G*_ represents s.d. of 4 experiments, while the colored shaded area of *V*_*D*_ curves presents s.e.m. of counted runs. (**C**) In the experiment, the positions of *V*_*D,i*_(*z*) *-V*_*G*_ intersections (circles, illustrated in B), together with the peaks (stars, illustrated in the insert figure) and the average positions (diamond) of bacteria density profiles all shift towards the front of the migration group for strains with the higher Tar-expression level, which has a higher chemotactic ability and migrate faster on agar plates (x-axis, see Method & Fig. S9). The related density profiles (PDF) were shown in the insert plot and the color-coding of lines/symbols in both panel C and D is the same as that in B. (**D**) The width of density profiles (2*σ*) of Tar-titrated bacteria decreases with the reversion rate *r*.

By comparing the statistics of cells with different Tar expression levels, we found that the expected drift velocity *V*_*D,i*_(*Z*) followed the same decreasing pattern from back to front (Fig 4B). More importantly, as the Tar-level (chemotactic ability) increases, the slope of the decreasing pattern increases, which is consistent to the model prediction shown in Fig 3A. The intersections between *V*_*D,i*_(*Z*) and *V*_*G*_, as well as the peak positions and mean positions of each tar-titrated density profiles (Fig 4C), shift toward the front as the chemotactic ability increases (measured by migration rate on agar plate (Cremer et al., 2019, Liu, Cremer et al., 2019)). The *V*_*D*_ cross point is always behind the peak position and the mean position (Fig 4C), suggesting that cells are leaking behind. Moreover, the width of each tar-titrated density profile (defined by 2*σ*_*i*_) decreases as the reversion rate r_i_ increases (Fig 4D), consistent with the model results in Fig 3C. Thus, as the O-U type model predicts, the width of the density profile is controlled by the reversion rate determined by the chemotactic ability *χ*_*i*_.

## Discussion

In summary, coordinated behaviors with ordered spatial arrangements of phenotypes are abundant in a wide range of biological and human-engineered systems and are believed to involved elaborate control mechanisms. For animal migrations, it is challenging to characterize simultaneously the computational strategy and behavior at individual levels so as to avoid averaging out phenotypic diversity, and the emergent behavior at population level (Couzin et al., 2005, Couzin et al., 2002, Vicsek & Zafeiris, 2012). Using bacterial chemotactic migration as a model system, we demonstrate that individual bacteria can spatially modulate their stochastic behaviors to perform mean reversion random motions around centers sequentially aligned by their chemotactic abilities, enabling a constant migration speed and ordered spatial arrangement of phenotypes at the collective level. Individual cells harness their own chemotactic system, together with collective consumption of attractant, to achieve the behavioral modulation, such that system transits from solitary to collective behaviors. This strategy of self-organization does not require sophisticated communications (Curatolo, Zhou et al., 2020, Karig, Martini et al., 2018, Liu, Fu et al., 2011, Payne, Li et al., 2013) nor other hydrodynamic interactions (Chen, Liu et al., 2017, Drescher et al., 2011, Zhang, Be’er et al., 2010) among individuals.

The behavior modulation depends on the chemotactic ability of individual, which is controlled by well determined chemotaxis related proteins. Amount them, we experimentally identified the abundance of the receptor protein Tar affects linearly the reversion rate and the width of dispersion. Simulation results suggests other key proteins that determines the basal tumble bias and the adaptation time may also affect the behavior modulation (Fig S8).

In the migratory group, the same rule of behavioral modulation applies to cells with different phenotypes, such that the random motions of cells are bounded by moving potential wells whose basin are sequentially aligned. However, it is noteworthy that cells could skip the potential wells from the back (Long, 2019), resulting leakage of cells in the migratory group (Holz & Chen, 1978, Novickcohen & Segel, 1984, Scribner, Segel et al., 1974). Phenotypes with weaker chemotactic abilities locating at the back of the group, where the effective potential well is shallower (Fig 3C), have more chance left to skip. Thus, such collective migration selects bacteria with higher chemotactic abilities (Liu et al., 2019).

The simple computational principle of behavioral modulation to allocate different phenotypes in the collective group is likely not limited to sensing the self-generated signal by consumption of attractant. Prominent example as trail-following migration (Couzin & Krause, 2003, Helbing, Keltsch et al., 1997), a typical class of collective behavior, a modified Langevin type model, where individuals tracing the accumulated signal secreted by all participants (Eq. S20), can reproduce similar spatiotemporal dynamics of behavioral modulation as well as ordered arrangements of phenotypes in the migratory group (Fig S10). Thus, this mechanism of matching individual abilities by the signal strength might provide an explanation of how other higher organisms organize ordered structures during group migration.

## Materials and Methods

### Strains

The wild type strain *Escherichia coli* (RP437) and its mutants used in this study were used in this study, where all plasmids were kindly provided by Dr. Chenli Liu. Specifically, the tar-titratable strain was constructed by recombineering according to a previous research (Zheng et al., 2016). Specifically, the DNA cassette of the *Ptet-tetR-tar* feedback loop was amplified and inserted into the chromosomal *attB* site by recombineering with the aid of plasmid *pSim5*. The *tar* gene at the native locus was seamlessly replaced with the *aph* gene by using the same recombineering protocol. To color-code the strains, we use plasmids with chloramphenicol resistant gene carrying YFP under constitutive promoter (for JCY1 strain) and *pLambda* drived mRFP1 plasmids maintained by kanamycin (For JCY2). To color-code tar-titratable strain (JCY20), a plasmid carrying YFP chloramphenicol resistant gene were transformed into constructedtar-titratable strain.

### Media and growth conditions

For bacterial culture, the M9 supplemented medium was used. The preparation of the M9 supplemented medium follows the recipe in previous study (Fu et al., 2018): 1×M9 salts, supplemented with 0.4% (v/v) glycerol, 0.1% (w/v) casamino acids, 1.0mM magnesium sulfate, and 0.05% (w/v) polyvinylpyrrolidone-40. 1×M9 salts were prepared to be 5×M9 salts stock solution: 33.9g · L^−1^ Na_2_HPO_4_, 15g · L^−1^ KH_2_PO_4_, 2.5g · L^−1^ NaCl, 5.0g · L^−1^ NH4Cl.

For migration experiments in the micro-channel, the M9 motility buffer was used. The recipe was: 1 × M9 salts, supplemented with 0.4% (v/v) glycerol, 1.0mM magnesium sulfate, and 0.05% (w/v) polyvinylpyrrolidone-40, 0.1mM EDTA, 0.01mM Methionine, and supplemented with 200μM aspartic acid.

For the migration rate measurements, the M9 amino acid medium with 0.2% (w/v) agar was used to prepare swim plate(Liu et al., 2019). The recipe was: 1 × M9 salts, supplemented with 0.4% (v/v) glycerol, 1 × animo acid, 200μM aspartic acid, 1.0mM magnesium sulfate, and 0.05% (w/v) polyvinylpyrrolidone-40. 1 × animo acid were prepared to be 5 × animo acid stock solution: 4mM alanine, 26mM arginine (HCl), 0.5mM cysteine (HCl · H_2_O), 3.3mM glutamic acid(K salt), 3mM glutamine, 4mM glycine, 1mM histidine (HCl · H_2_O), 2mM isoleucine, 4mM leucine, 2mM lysine, 1mM methionine, 2mM phenylalanine, 2mM proline, 2mM threonine, 0.5mM tryptophane, 1mM tyrosine, 3mM valine. All experiments were carried out at 30 °C. Plasmids were maintained by 50 μg · mL^−1^ kanamycin or 25 μg · mL^−1^ chloramphenicol.

### Sample preparation

The bacteria from frozen stock was streaked onto the standard Luria-Bertani (LB) agar plate with 2% (w/v) agar and cultured at 37°C overnight. 3-5 separate colonies were picked and inoculated in 2mL M9 supplemented medium for overnight culture with corresponded antibiotics to maintain plasmids. The overnight culture was diluted by 1:100 into 2mL M9 supplemented medium the next morning. For Tar titration strains, related aTc were added in this step. When the culture OD600 reaches 0.2-0.25, it was then diluted into pre-warmed 15mL M9 supplemented medium so that the final OD600 was about 0.05 (Liu et al., 2019, Zheng, Bai et al., 2020, Zheng et al., 2016).

Bacteria were washed with the M9 motility buffer and were re-suspended in fresh M9 motility buffer to concentrate cell density at OD600 about 1.0. Then, the wild type strain and fluorescent strain were mixed with ratio of 400:1 before loaded in the microfluidic chamber (Fu et al., 2018, Saragosti et al., 2011). For Tar titration experiments, the wild type strain (RP437) was mixed with two fluorescent strains (JCY2 & JCY20) by 400:8:1.

### Microfabrication

The microfluidic devices were fabricated with the same protocol and the same design as previous research (Bai, Gao et al., 2018, Fu et al., 2018), except that the capillary channel was designed longer than that of previous ones. The size of the main channel was 20mm × 0.6mm × 0.02mm and only one gate at the end of the channel was kept (Fig S1A).

### Band formation

Sample of mixed cells with density OD600 ≈ 1.0 was gently loaded into the microfluidic device and then the device was spun for 15min at 3000*rpm* in an 30 °C environmental room so that almost 1~1.5 · 10^5^ cells were placed to the end of the channel. After spinning, the microfluidic device was placed on an inverted microscope (Nikon Ti-E) equipped with a custom environmental chamber set to 50% humidity and 30 °C.

### Imaging

The microscope and its automated stage were controlled by a custom MATLAB script via the μManager interface (Edelstein, Tsuchida et al., 2014, Fu et al., 2018). A 4X objective (Nikon CFI Plan Fluor DL4X F, N.A. 0.13, W.D. 16.4 mm, PhL) was placed in the wave front and the fluorescent bacteria, seen as randomly picked samples of the migrating group, were captured continuously in 10 mins until they leave the view. Time-lapsed images with YFP fluorescence of the migrating cells were acquired by a ZYLA 4.2MP Plus CL10 camera (2048 × 2048 array of 6.5 × 6.5 *μm* pixels) at 9 frames/s (fps) through. A LED illuminator (0034R-X-Cite 110LED) and an EYFP block (Chroma 49003; Ex: ET500/X 20, Em: ET535/30 m) compose the lightening system.

For the Tar titration experiments, the channel was first scanned with 10X objective (CFI Plan Fluor DL 10X A, N.A. 0.30, W.D. 15.2mm, PH-1) enlighten by a LED illuminator (0034R-X-Cite 110LED) through the RFP block (Chroma 49005, Ex: ET545/X 30, Em: ET620/60 m) and EYFP block channels for 7 neighbored views around the migration group. These images were further combined to 2 large pictures of the RFP fluorescent strains and YFP fluorescent strains. The channel was scanned twice, respectively before and after the 10 mins tracking of fluorescent Tar titrated cells.

### Tracks extraction and state assignment

The acquired movie was first analyzed with the U-track software package to identify bacteria and to get their trajectories (Jaqaman, Loerke et al., 2008). Then the tracks were labeled by run state and tumble state by a custom MATLAB package (Waite et al., 2016) using a previously described clustering algorithm (Dufour et al., 2016).

### Track analysis

The group velocity V_G_ was calculated by averaging the frame to frame velocity (*dt* ≈ 0.11*s*) over all tracks and all time. The cell number for the first frame over a spatial bin of Δx = 60μm and a channel section *a* = 12000*μm*^2^ were calculated to get the density profile 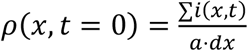. The peak position of the first frame (*x*_*peak*_(*t* = 0)) was then determined by the maximum of *ρ*(*x, t* = 0). The position of each bacterium (*x*_*i*_(*t*)) was transformed to moving coordinate position z_i_ by the group velocity *V*_*G*_ and origin of the axis on the density peak by *z*_*i*_ = *x*_*i*_(*t*) − *V*_*G*_ *t* − *x*_*peak*_(*t* = 0). Given the relative position of each cell, we recalculated the density profile in moving coordinate 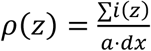. The width of the density profile was defined by two times the standard deviation of relative positions 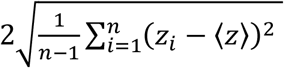. The spatial distribution of the instantaneous velocities ⟨*V*_*I*_ (*z*)⟩ were calculated by averaging the velocity in spatial bin of *Δz* = 240*μm*.

A tumble-run event is the minimal element of bacterial behavior. The typical spatial scale of a tumble-run event is about 20μm, which is much smaller than the spatial bin size chosen in this study (240μm). The spatial distributions of run time ⟨*τ*_*R*_(*z*)⟩, tumble time ⟨*τ*_*T*_(*z*)⟩ and run length ⟨*l*_*R*_(*z*)⟩ were calculated by averaging the related values of all the events with tumbling position (*z*_*T*_) located in each spatial bin (*z*). As the displacement of tumble is small, the tumbling position is approximately the starting position of runs. For each tumble-run event, we have the vector linking starting position and end position of the run. The running angle *θ*_*R*_ is then defined by the angle between run direction and the group migration direction. One can easily deduce all the other quantities with the formulations in Table 1.

**Table 1.** Summary of quantities.

| Quantities | Definition | Formulations |
| --- | --- | --- |
| $V_G$ | Group velocity | $V_G = \langle \frac{dx}{dt} \rangle$ |
| $z$ | Moving coordinate | $z = x - V_G t$ |
| $V_I(z)$ | Instantaneous velocity | $V_I(z) = \langle \frac{dx(z)}{dt} \rangle$ |
| $B_\tau(z)$ | Run time bias | $B_\tau(z) = \frac{\langle \tau_R(z) \cdot \cos \theta_R(z) \rangle}{\langle \tau_R(z) \rangle}$ |
| $B_l(z)$ | Run length bias | $B_l(z) = \frac{\langle l_R(z) \cdot \cos \theta_R(z) \rangle}{\langle l_R(z) \rangle}$ |
| $V_D(z)$ | Expected drift velocity | $V_D(z) = \frac{\langle L_R(z) \cdot \cos \theta_R(z) \rangle}{\langle \tau_R(z) + \tau_T(z) \rangle}$ |
| $\rho(z)$ | Cell density | $\rho(z) = \frac{\sum i(z)}{a \cdot \Delta z}$ |
| $S(z)$ | Chemo-attractant concentration | $-V_G \frac{dS}{dz} = D_s \frac{\partial^2 S}{\partial z^2} - k\rho$ |
| $g(z)$ | Perceived gradient | $g(z) = \frac{d \ln \left( \frac{1+S(z)/K_{off}}{1+S(z)/K_{on}} \right)}{dz}$ |

### Growth rate and migration rate measurement

Growth rates of Tar-titrated strains were calculated from exponential fitting (*R*^2^ > 0.99) over measured curves of cell density (OD600) with respect to time. A 250 mL flask with 20 mL M9 supplement medium were used. All measurements were performed in a vibrator of rotation rate of 150 rpm at 30°C. OD600 was measured by a spectrophotometer reader every 25 min. Each strain has been measured for at least three times.

The semi-solid agar plate was illuminated from bottom by a circular white LED array with a light box as described previously (Liu et al., 2011, Liu et al., 2019, Wolfe & Berg, 1989) and was imaged at each 2 hours by a camera located on the top. As bacteria swimming in the plate forms ‘Adler ring’, we used the first maximal cell density from the edge to define the moving edge of bacterial chemotaxis. The migration rate was then calculated from a linear fit over the data of edge positions in respect to time (*R*^2^ > 0.99).

### Models and simulations

Details of the theoretical models and numerical simulations were presented in the appendix notes. In which, the Langevin equation was deduced and solved numerically with a particle-based simulation; the approximated OU type equation and its traveling wave solution was deduced; an agent-based simulation of bacterial with chemotaxis pathway was performed.

## Acknowledgments

The authors acknowledge C.Liu, for sharing E.coli strains and plasmids; S. Huang for help with the microfluidics; T. Emonet for help with single cell tracking and agent-based simulations; T. Hwa, F. Jin, L. Luo, J. Long for insightful discussions.This work was financially supported by the National Key R&D Program of China (2018YFA0903400), the National Natural Science Foundation of China (32071417), CAS Interdisciplinary Innovation Team (JCTD-2019-16), Guangdong Provincial Key Laboratory of Synthetic Genomics (2019B030301006), Shenzhen Peacock Plan (KQTD2016112915000294) to X.F., the National Natural Science Foundation of China (11804355, 31770111), the Guangdong Provincial Natural Science Foundation (2018A030310010), Shenzhen Grant (JCYJ20170413153329565) to Y.B..

## Author contributions

X.F. initiated and directed the research. C.H. constructed strains, fabricated micro-fluidics and tracked bacterial motions. Y.B. analyzed data, performed the particle-based simulations and the agent-based simulations. Y.B., X.L., and X.F. analyzed the results and mathematical modelling. All authors contribute in writing the manuscript.

## Data and materials availability

All data needed to evaluate the conclusions in the paper are present in the paper and/or the Supplementary Materials. Codes for bacterial tracking and agent-based simulation were presented in previous papers and were modified with details presented in the Appendix. The OU model were simulated with Matlab and related codes were avablable on line: https://github.com/BaiYangBqdq/spatial_modulation_in_group_migration

